# On the Prediction of non-CG DNA Methylation

**DOI:** 10.1101/2022.04.26.489600

**Authors:** Saleh Sereshki, Michalis Omirou, Dionysia Fasoula, Stefano Lonardi

**Affiliations:** Department of Computer Science and Engineering, University of California Riverside, Riverside, CA 92521; Agricultural Microbiology Laboratory, Agricultural Research Institute, Nicosia, 1516 Cyprus; Department of Plant Breeding, Agricultural Research Institute, Nicosia, 1516 Cyprus

## Abstract

DNA cytosine methylation is an epigenetic modification that has a critical role in gene regulation and genome stability. DNA methylation can be detected and measured using sequencing instruments after sodium bisulfite conversion, but experiments can be expensive for large eukaryotic genomes. Sequencing non-uniformity and mapping biases can leave parts of the genome with low or no coverage, thus hampering the ability of obtaining DNA methylation levels for all cytosines. To address these limitations, several computational methods have been proposed that can predict DNA methylation from the DNA sequence around the cytosine, or from the methylation level of nearby cytosines. Most of these methods are, however, entirely focused on CG methylation in humans and other mammals. In this work, we study for the first time the problem of predicting cytosine methylation for CG, CHG, and CHH contexts on five plant species, either from the DNA primary sequence around the cytosine or the methylation levels of neighboring cytosines. In this framework, we also study (1) the cross-species prediction problem, i.e., the classification performance when training on one species and testing on another species, and the (2) the cross-context prediction problem, i.e., the classification performance when training on one context and testing on another context (within the same species). Finally, we show that providing the classifier with gene annotation information allows our classifier to outperform the prediction accuracy of state-of-the-art methods.

## 1 Introduction

DNA methylation is an epigenetic mark that has a critical role in regulating a variety of cellular processes, e.g., gene expression, genome stability, and gene imprinting (see, e.g., [1, 2, 3]). The most common type of DNA methylation is the addition of a methyl group to the fifth carbon of a cytosine residue. In mammals, DNA methylation is mostly found at cytosines that are followed by guanine base, known as CG methylation. Long stretches of DNA that are very rich in the dinucleotide CG, called CpG islands, tend to be less methylated than the other cytosines in the genome [4, 5, 6]. As said, DNA methylation is one of several epigenetic mechanisms that cells use to regulate gene expression [7, 8]. In humans, the dysregulation of DNA methylation is associated with a variety of diseases (e.g., cancer [9, 10]) and neurological disorders (see, e.g., [11, 12]). In plants and other non-vertebrates, cytosine methylation in the CHH and CHG contexts (where H represents any base except G) is almost as common as methylation in CG context [13, 14]. It is now well-understood that distinct molecular mechanisms in the cells regulate cytosine methylation and demethylation depending on the context [15, 16, 17, 18].

Recent studies suggest the importance of non-CG methylation in both vertebrates and non-vertebrates. In humans, non-CG methylation is the most abundant form of DNA methylation in neurons and plays a critical role in cognitive functions (see, e.g., [19, 20, 21]). Dysregulation of type of methylation has been associated with mental diseases like schizophrenia [22]. In plants, it has been shown that (i) distinct pathways and molecular processes maintain cytosine methylation in CG, CHG, and CHH contexts (see, e.g., [15, 23]), (ii) gene body methylation patterns differ for CG and non-CG methylation (see, e.g., [24, 23, 25, 26]), (iii) both CG and CHG methylation are correlated to genome size and repetitive content, while CHH methylation is not [27].

Several methods are available for reading the methylation status of cytosines. Whole genome bisulfite sequencing (WGBS or BS-Seq) is arguably the most common. Other techniques include bead chip arrays (e.g., Illumina Infinium), Oxford Nanopore [28], or affinity enrichment-based, like methylcytosine-specific antibodies (MeDIP-seq). BS-Seq allows for quantitative cytosine methylation detection at single-base resolution and is still considered the “gold standard” for the analysis of DNA methylation. By treating DNA with sodium bisulfite, unmethylated cytosines transform to uracils, while methylated cytosines stay intact. Once the DNA is converted, DNA sequencing (typically carried out on Illumina instruments) generates the reads which are then mapped to the reference genome using conversion-aware mapping tools (e.g., Bismark [29], BS Seeker [30] or BRAT-nova [31]). Since the methylation level for each cytosine is obtained by computing the ratio between the number of reads that indicates a methylated cytosine and the total number of reads, the statistical confidence associated with this measurement depends on the depth of sequencing coverage at each cytosine. To guarantee that the read coverage is sufficiently for all the cytosines in the genome, the average sequencing depth needs to be high, which can be expensive for large eukaryotic genomes. Since sequencing depth is not uniform across the genome, some cytosines can end up with low or no read coverage, which prevents the accurate measurement of their methylation level. This problem is particularly acute for single-cell experiments because the coverage is usually much lower and less uniform than bulk sequencing data.

As a result, several methods have been developed in the last ten years for predicting or imputing cytosine methylation levels. These methods are mostly focused on prediction of CG methylation in humans. In one of these studies, a deep neural network uses sequence and methylation level of neighboring cytosines to predict the methylation level from single-cell experiments, exclusively for the CG context in human and mouse cells [32]. Another method targeted at CG methylation prediction on mouse single-cell data is proposed by Li *et al*. [33] using again deep learning. Their model uses the underlying DNA sequence, the methylation status and distance of neighboring cytosines to carry out methylation prediction. In [34], a convolutional neural network is proposed for the prediction of CG methylation levels from the DNA sequence in the human genome. In [35], the authors propose a convolutional neural network for predicting the cytosine methylation in the CG context from the DNA sequence data in humans. Recently, Waele *et al*. use transformer architecture for imputation of single-cell methylation levels in human and mouse [36].

Other studies use Illumina Infinium Human Methylation 450 array data to carry out predictions. In one of these studies, Zhang et al. [37] propose a random forest model that uses the DNA sequence, the neighboring cytosines methylation levels and the presence of CpG islands to predict CG methylation levels in humans. In a similar study, the authors of [38] use a random forest model to predict cytosine methylation levels in humans from Infinium methylation levels and the distance of neighboring CG.

All these studies demonstrate that it is possible to predict CG methylation from the DNA sequence or the neighboring methylation levels at various levels of accuracy. However, the problem of predicting non-CG methylation has been so far largely ignored despite its growing importance in molecular biology. Even worse, sometimes non-CG methylation is improperly bundled with CG prediction, despite clear mechanistic differences at the cellular level. Here we address for the first time, to the best of our knowledge, this fundamental shortcoming. Specifically, our work makes the following contributions. (1) We study the problem of predicting cytosine methylation independently for the CG, CHG, and CHH context (and for all three contexts mixed) on five plant species on either the DNA primary sequence or the methylation level of neighboring cytosines. (2) We study the cross-context prediction problem, i.e., we investigate how hard is to predict methylation for a specific context when trained on a different one. (3) We study the cross-species prediction problem, i.e., we investigate how hard is to predict methylation for a specific species when trained on a different one. (4) We show that one can obtain higher predictive accuracy from the levels of neighboring cytosines than from the DNA sequence. (5) We show that providing gene annotations allows our prediction method to outperform state-of-the-art methylation prediction methods.

## 2 Results

As said earlier, while vertebrate DNA cytosine methylation is primarily found in the CG context, plants have significant levels of DNA methylation in the CG, CHG and CHH contexts [39, 15]. To focus on CHG and CHH methylation we selected five plant species for which BS-Seq data was available, namely (1) *Arabidopsis thaliana* representing the Brassicales order, (2) rice (*Oryza sativa*) representing the Paoles order (the only monocotyledons in our study), (3) tomato (*Solanum lycopersicum*) representing the Solanales order, (4) cucumber (*Cucumis sativus*) representing the Cucurbitales order, and (5) cowpea (*Vigna unguiculata*) representing the Fabales order. Data sources and processing of BS-Seq reads are described below in Methods. Supplementary Table 2 summarizes the main statistics of the BS-Seq reads for each plant species.

The average cytosine coverage from BS-Seq mapped reads ranged from 5x in tomato to 21x in Arabidopsis (see Supplementary Table 3). To ensure sufficient statistical confidence in the determination of methylation levels, a strict threshold for coverage was adopted; we only called cytosines that were covered by more than ten reads. A cytosine was considered methylated if more than half of the reads covering it indicated methylation (and unmethylated otherwise). Observe that the percentage of CG, CHG and CHH methylation varies greatly among different species. Supplementary Table 4 shows that more than 89% of cytosines in the CG context are methylated in tomato compared to only about 27% in Arabidopsis; more than 62% of cytosines in the CHG context are methylated in tomato compared to only 11% in Arabidopsis; 2.73% of cytosines in the CHH context are methylated in tomato compared to 0.68% in Arabidopsis.

Our cytosine prediction analyses can be logically organized in five steps.

In the first step, context-specific species-specific data sets were used to train a simple Random Forest (RF) classifier, which has been used frequently in the literature for cytosine methylation prediction (see, e.g., [37, 38]). The input to RF was a sequence of 3,200 nucleotides (hereafter called *window size*) centered at the cytosine under consideration and labeled with the binary methylation status of the center cytosine. The training set was composed by 500 thousand DNA sequences selected uniformly at random from the genome (for each specific methylation context), where half of the sequences had a center cytosine which was methylated, and half of the sequences had a center cytosine which was unmethylated. The training set was balanced because the highly skewed distribution in some contexts could make the prediction trivial. For example, more than 99% of cytosines in the CHH context for Arabidopsis are unmethylated, thus a “classifier” that predicts every cytosine in the CHH context for Arabidopsis to be unmethylated would achieve more than 99% accuracy. In contrast, almost 90% of cytosines in the CG context for tomato are methylated, thus a “classifier” that predicts every cytosine in the CG context for tomato to be methylated would achieve almost 90% accuracy. RF was also trained on all contexts mixed in equal proportions. For the combined context (indicated as “ALL” in the figures), we balanced the three contexts in equal proportions because otherwise the skewed distribution in some contexts could make the prediction trivial. For example, for tomato a classifier that calls (i) all cytosines in the CG context methylated, (ii) all cytosines in the CHG context methylated, and (iii) all cytosines in the CHH context unmethylated would achieve an expected 92% accuracy, based on Supplementary Table 6 and some basic probability calculations (not shown).

Figure 1-A shows the cross-validation accuracy of RF on Arabidopsis, cowpea, rice, cucumber, and tomato, independently for each context (CG, CHG, CHH) and for all contexts mixed. First, observe that the prediction performance of cytosine methylation from the sequence is highly dependent on the context. Also observe that the prediction of methylation in the CHH context is often more accurate than the prediction of methylation in the other two contexts, which suggests that the CHH methylation could be more sequence-dependent in plants than CG or CHG. Mixing all the contexts results in a decrease in classification performance, which supports the need of an individual classifier for each context.

**Figure 1:**
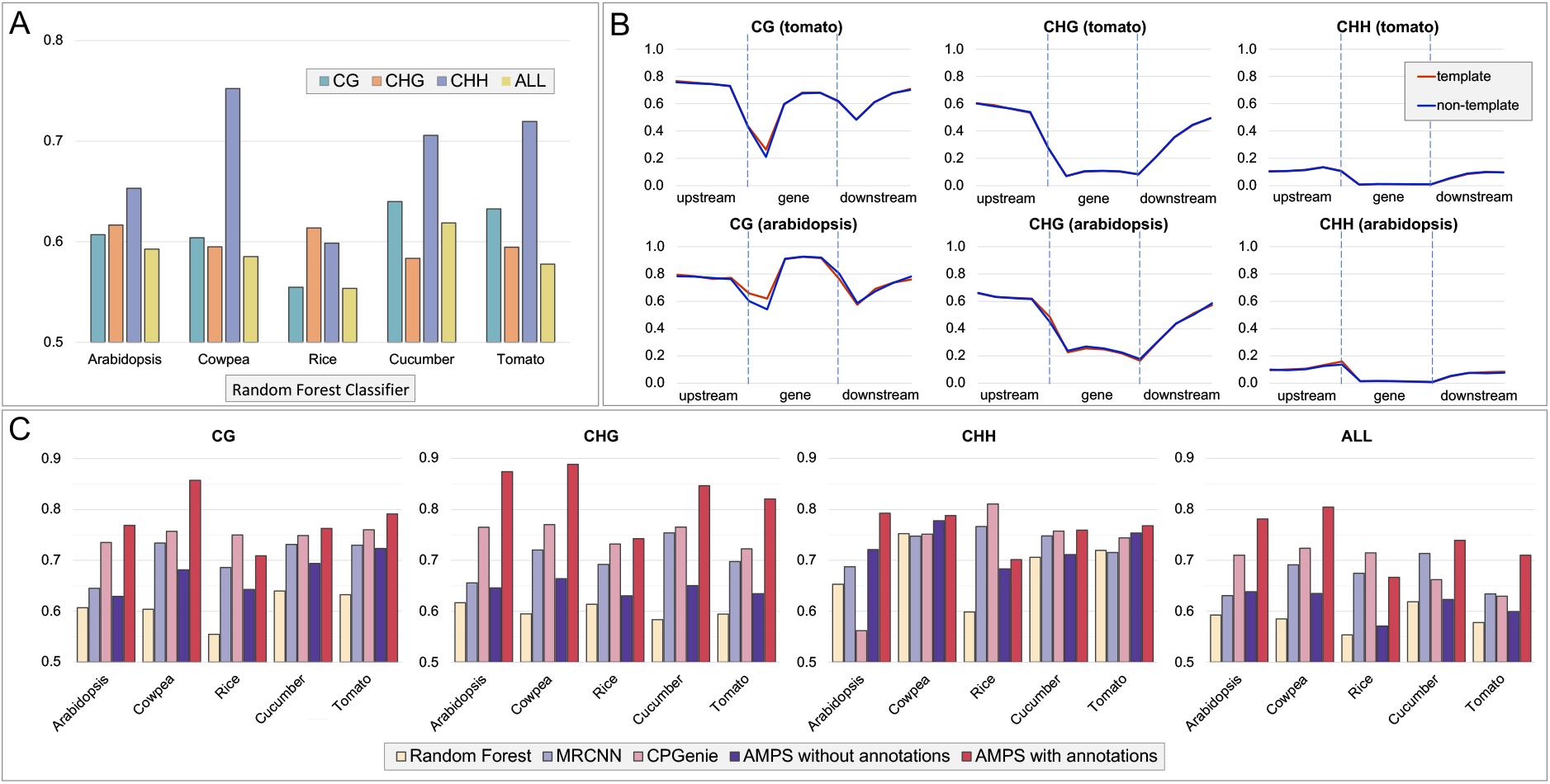
**A:** Context-specific species-specific cross-validation prediction accuracy of a simple Random Forest classifier on the five plant species included in this study; **B:** context-specific gene body methylation levels for tomato (top row) and Arabidopsis (bottom row) for the template and non-template strand; **C:** Context-specific species-specific cross-validation prediction accuracy for Random Forests, MRCNN, CPGenie, AMPS without annotations and AMPS with annotations (AMPS is the new method proposed here).

In the second step, we investigated the observations reported in the literature that the methylation levels in Arabidopsis and other plant species varied drastically in gene bodies compared to upstream and downstream regions [24, 23, 25, 26]. Figure 1-B shows the methylation levels for template and non-template strands in the gene body and 2 Kb flanking regions averaged over all genes in tomato and Arabidopsis (see Methods for details). Supplemental Figure 1 shows the corresponding analyses for the other plant species in this study. Observe that CG and CHG methylation levels dip in correspondence to the gene boundaries, and that overall shape of methylation levels is context dependent.

These analyses prompted the question on whether providing the classifier with genomic annotation information (e.g., gene boundaries, coding sequence boundaries, intron/exon boundaries) could boost the classification performance for cytosine methylation. To answer this question, we designed a new classifier that uses the annotations listed above in addition to the DNA sequence. Our classifier, called <u>A</u>nnotation-based <u>M</u>ethylation <u>P</u>rediction from <u>S</u>equence (AMPS), is simple deep learning architecture that uses convolutional neural networks (details in the Methods section). Since we planned to use annotations related to genes and introns, we investigated how much of each genome is annotated by these genomic features Supplemental Figure 7 shows that the 65% of the smallest genome (Arabidopsis) is annotated as a gene, while only 17% of the largest genome (tomato) is annotated as a gene. Supplemental Figure 8 shows the context-specific species-specific fraction of all cytosines covered by annotations. To determine whether annotations would improve the classification accuracy, we carried out a comparative analysis with previously published methods which led to the third step in the analysis.

In this step, we compared the performance of AMPS to Random Forests, CpGenie [35] and MRCNN [34]. We chose to compare AMPS against CpGenie and MRCNN because they are considered state-of-the-art methods for methylation prediction exclusively from DNA sequence. In fairness, we should note that CpGenie and MRCNN were optimized for predicting methylation in the CG context on the human genome. We retrained CpGenie and MRCNN on our species-specific and context-specific data set, but their architectures might not be optimal for non-CG non-human methylation. We should also note that CpGenie uses a more sophisticated deep-learning architecture than AMPS, resulting in a larger number of weights and hyper-parameters. MRCNN has four convolution layers followed by two fully connected layers while AMPS has two convolution layers followed by one fully connected layer. The hyper-parameters of the AMPS were not highly optimized to ensure that the method would be able to generalize, but the effect of window size and the training set size on the prediction performance was extensively studied in Section 2.3. In all experiments, AMPS’ window size (with or without annotation) was 3.2 Kb, CpGenie’s window size was 1 Kb, and MRCNN’s window size was 400 bp. These two latter window sizes were dictated by the corresponding architectures proposed by the authors. All classifiers were trained on 500 thousand DNA sequences selected uniformly at random from the genome (a discussion about training set size can be found in Methods). As explained in the Section ‘Training Set Design’, the variance in performance across multiple random samples was negligible, so all the experiments were carried out on a single sample to reduce the overall computational cost.

Figure 1-C reports the accuracy of the classifiers listed above, including AMPS without annotations. Observe that (i) on the CG, CHH and ALL contexts, AMPS (with annotations) outperformed the other methods in four out of the five species (on rice, CpGenie performed better), (ii) on the CHG context, AMPS (with annotations) outperformed the other methods on all species, (iii) AMPS (with annotations) had the biggest improvement over AMPS without annotations on the CHG context (which is the context in Supplemental Figure 8 that has the highest percentage of cytosines covered by annotations, irrespective on the species), (iv) in some cases, the accuracy of AMPS without annotation was lower than other predictors, suggesting the critical advantage of using genomics annotation as an input feature. Also, observe in Figure 1-C that (i) the accuracy of different classifiers is context dependent and (ii) in 20 out of 25 experiments the prediction accuracy that used all the contexts mixed was lower than training on each context independently. The same experimental results are shown in Supplementary Figure 4 but grouped by classifier instead of species.

### 2.1 Cross-context and cross-species prediction

In the fourth step, we investigated the ability of the predictor to carry out cross-species prediction from the DNA sequence. Figure 2-A shows the accuracy of AMPS (without annotations) when trained with one species and tested on another, for each context individually and all contexts mixed. Annotations could not be used for this task because of the different number of functional elements available for each organism (e.g., three for Arabidopsis, and two for cowpea). Observe that training and testing on the same species achieves the highest accuracy, as expected.

**Figure 2:**
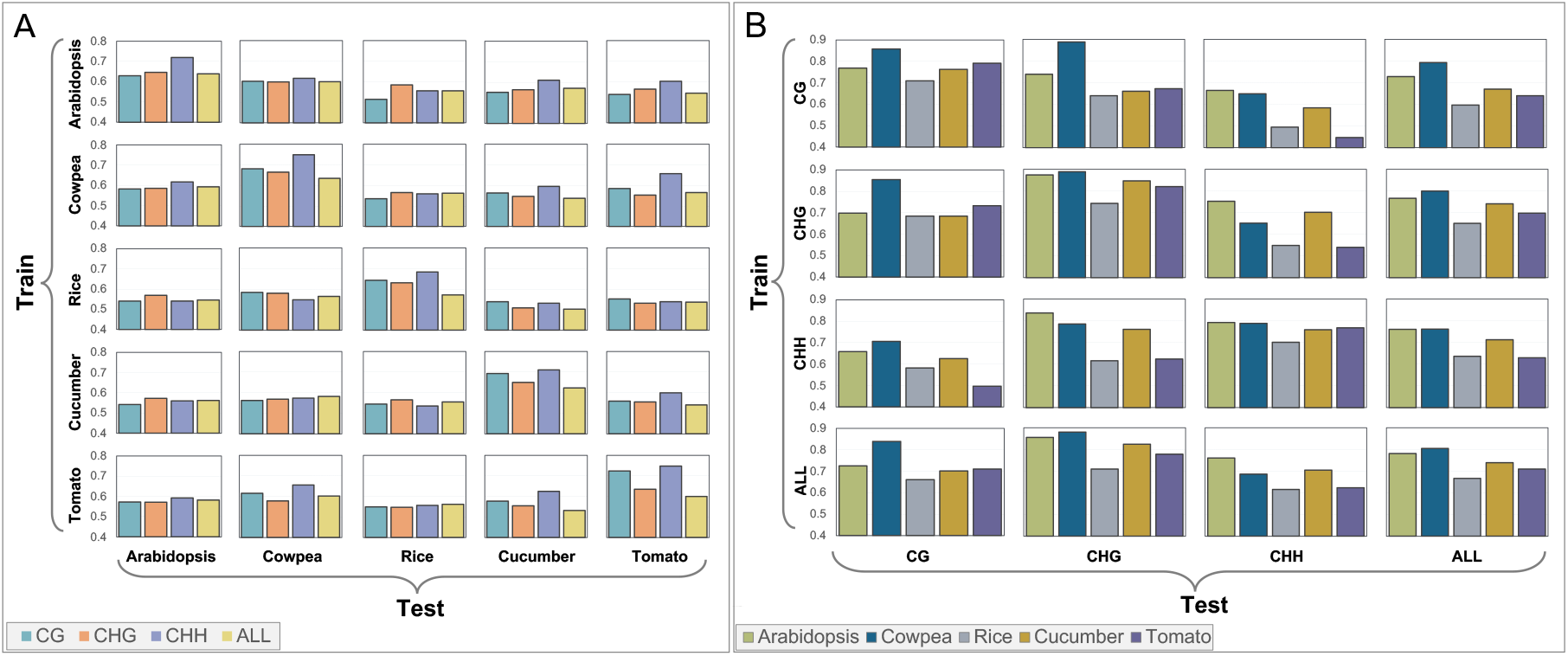
**A:** Cross-species methylation prediction accuracy of AMPS without annotation; **B:** Cross-context methylation prediction accuracy of AMPS with annotations.

In the fifth step, we investigated cross-context predictions. The performance of AMPS (with annotations) was evaluated when trained with one context and tested on another. Figure 2-B shows the prediction accuracy for all pairs of training/testing contexts (including the mixed contexts) for all species. Observe that in the most of the cases, cross-context prediction accuracy is the highest when training and testing on the same context, as expected, but not when all contexts are mixed. A similar observation can be made on the cross-context prediction accuracy for the AMPS without annotation (Supplementary Figure 5).

### 2.2 Prediction based on neighboring cytosines

In this section, we studied the problem of predicting cytosine methylation from the methylation levels of the neighboring cytosines, which is common in literature for data imputation. In this case, the classifier took in input a vector of methylation levels (half upstream and half downstream, with the condition that cytosines had to have a sufficient read coverage to be included) and predicted the binary methylation status of the center cytosine. In all experiments, the size of the vector was twenty components. The data set size was fifty thousand methylation vectors uniformly sampled from the genome, in which half of them were centered at a methylated cytosine while the other half was centered at an unmethylated cytosine. Eighty percent of the data set was used for training, ten percent was used for validation, and ten percent was used for testing. Our classifier was a simple fully connected neural network with four hidden layers (more details in Methods). As we did earlier, we carried out methylation prediction for each species and for each context individually, but also for all contexts combined. Figure 3-A shows that the prediction accuracy is again context- and species-specific. More specifically, observe that (i) cytosine methylation in the CG context is the easiest to predict, while methylation in the CHH context is the hardest (somewhat the opposite of what we observed for sequence-based prediction, as shown in Figure 1), (ii) combining the contexts degrades the prediction performance compared to context-specific classifiers, (iii) methylation prediction in tomato appears to be harder than other species.

**Figure 3:**
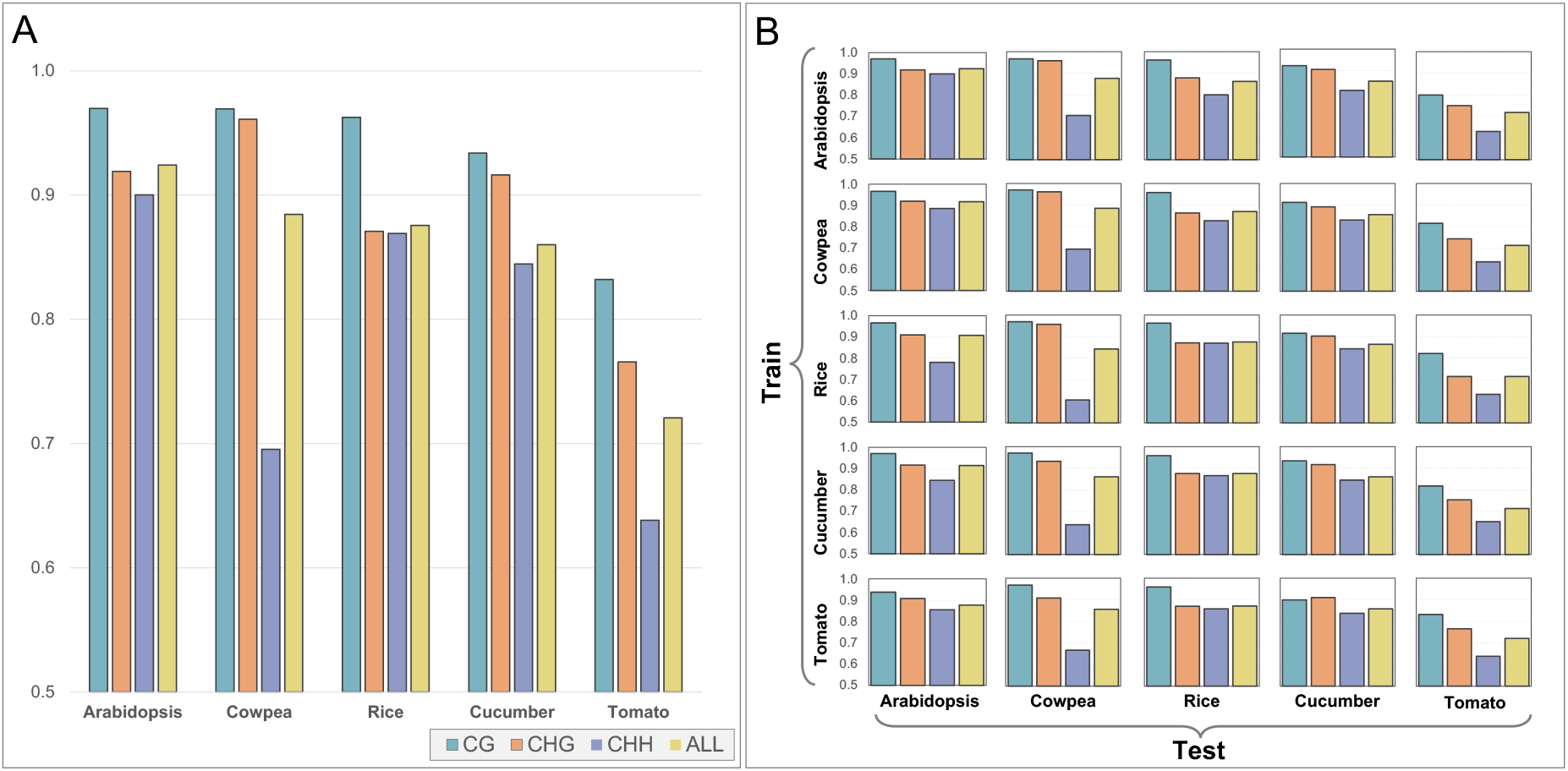
**A:** Methylation prediction accuracy from the methylation levels of the neighbouring cytosines; **B:** Cross-species methylation prediction accuracy from the methylation levels of the neighbouring cytosines

We also carried out cross-species and cross-context predictions using neighboring cytosines. Figure 3-B shows the prediction accuracy when training on a species and testing on another. Observe that the accuracy does not change significantly as one moves down the rows of the matrix. That indicates that the prediction accuracy is somewhat independent from the species the predictor was trained on, which is again different from what we observed for sequence-based prediction (as shown in Figure 2-A). Also observe that the accuracy appears to be correlated with the average cytosine coverage (Supplementary Table 3). For instance, the worst overall accuracy is for tomato, which has the lowest average cytosine coverage. The best overall accuracy is for Arabidopsis, which has the highest average cytosine coverage. Supplemental Figure 6 shows the prediction accuracy when training on a context and testing on another. With some exceptions, observe again that the accuracy does not significantly change as one moves down the rows. This implies that the prediction accuracy is somewhat independent from the context on which the classifier was trained on.

### 2.3 Effect of the window size and training set size on the prediction accuracy

Two critical parameters for the prediction accuracy from the DNA sequence are (i) the size of the training set and (ii) the size of the input sequence (or *window size*). Here we carried out extensive tests to determine the optimal values for these two parameters using AMPS (with annotations) as a classifier.

As expected, the size of the training set directly affects the performance of the classifier. We recorded the accuracy of AMPS (with annotations) on all species and all contexts using a data set with 40K, 80K, 120k, 200K, 400K, 600K, 800k, and 1M sequences. Eighty percent of the data was used for training, ten percent was used for validation and ten percent was used for testing. For some (organism, context) pairs, the number of cytosines that had sufficient read coverage to be called methylated (or not) was insufficient to satisfy the data set needs. In those cases, the larger data sets are missing from the analysis and the figures.

Figure 4-A illustrates AMPS’ accuracy as function of the data set size for cowpea, while Supplementary Figure 2 shows the corresponding plots for all plant species. For Arabidopsis, we trained/tested AMPS six times. Supplemental Figure 2 shows the standard deviation of these six experiments as error bars. Observe that the standard deviations are very low, which justifies the time-saving choice to rely on one experiment for all other tests. Also, observe that for data set with 400 thousand sequences or more, the accuracy is high and relatively stable in all plant species. Based on this observation we used data sets composed of 500 thousand sequences, if there were sufficient cytosines available. If there were not, we used all the available cytosines.

**Figure 4:**
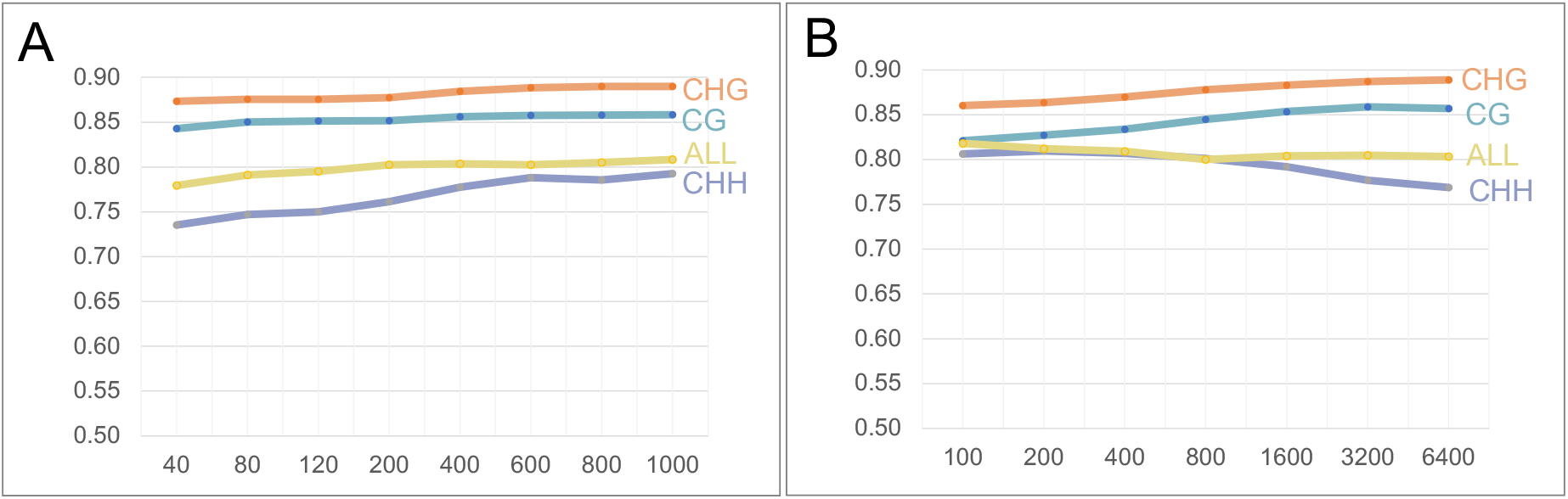
A. Prediction accuracy as a function of the training set size for AMPS with annotations (the x-axis represents thousands of samples) on the cowpea dataset; B. Prediction accuracy as a function of the window size for AMPS with annotations (the x-axis represents the window size in base pairs) on the cowpea dataset

Figure 4-B shows AMPS’ accuracy as function of the window size (100 bp, 200 bp, 400 bp, 800 bp, 1600 bp, 3200 bp and 6400 bp) on the cowpea genome. Supplemental Figure 3 shows the corresponding plots for other species. As we did earlier, we trained/tested AMPS six times on Arabidopsis. Supplemental Figure 3 shows the standard deviation of these six experiments as error bars. Observe that the standard deviations are still very low. When comparing the graphs in Supplemental Figure 3, observe that context-specific predictions are differently affected by the window size. For CG and CHG, the prediction accuracy increases up to a window size of 3,200 bp. But for CHH the accuracy does not change or degrade by increasing the window size. We do not have an explanation for this phenomenon.

## 3 Methods

### Data sources and data pre-processing

Whole Genome Bisulfite Sequencing (BS-Seq) data for *Arabidopsis thaliana*, rice (*Oryza sativa*), tomato (*Solanum lycopersicum*) and cucumber (*Cucumis sativus*) were obtained from the Sequence Read Archive (SRA) of NCBI/NIH. BS-Seq data for cowpea (*Vigna unguiculata*) were generated in the context of the Cyprus national project “Cowpea breeding and adaptation to climate change” [40]. The cowpea genome was recently sequenced and assembled by our group [41]. The other genomes were obtained from NCBI (see Supplementary Table 1 for source and assembly versions).

Supplementary Figure 9 shows the location of these five species on a phylogenetic tree of the major land plant species [42]. Observe that these plant species belong to five distinct orders: Arabidopsis belongs to the Brassicales, cowpea to the Fabales, cucumber to the Cucurbitales, tomato to Solanales, and rice to the Poales. Not all plant orders are represented in our study, but we plan to expand it to the other orders in the future.

Read quality was checked using FastQC v0.11.5. In some cases, sequencing primers were detected in the sequenced reads. Reads that had these anomalies were trimmed with Trimmomatic v0.33 [43]. Reads were mapped against the corresponding reference genome using Bismark v0.22.2 using default parameters [29]. Only reads that were uniquely aligned were used by Bismark, i.e., ambiguous reads with multiple mappings were discarded.

The output of Bismark was processed using custom scripts as follows. First, the methylation level of each cytosine was obtained by computing the ratio of the number of methylated reads over all the reads covering that cytosine. A cytosine was declared to be *methylated* if the methylation level was at least 0.5, *unmethylated* otherwise. Methylation was only called for cytosines that had a coverage of more than ten reads. Cytosines covered by less than eleven reads had an unknown methylation status and were excluded from all experiments.

### Gene body methylation profiles

To obtain the average species-specific gene methylation profile, we collected the methylation levels for each annotated gene, as well as the methylation levels in 2000 base pairs upstream and downstream of each gene. Gene bodies and flanking regions were split into five bins each, for a total of 15 bins. For each bin *r* ∈ [1, 15] the average methylation level *M* (*r*) was calculated as follows

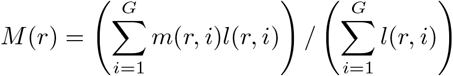

where *G* is the total number of annotated genes in that species, *m*(*r, i*) is the ratio of methylated cytosines over all cytosines in bin *r* of gene *i*, and *l*(*r, i*) is the length of bin *r* in gene *i* (the length of a bin is 400 base pair for flanking regions, while it is equal to one fifth of each gene length for the bins within a gene).

### Training set design

For classifiers that rely on the DNA sequence, a context-specific training set was composed by *n* DNA sequences of length *W*_*s*_ centered at a cytosines (i.e., *W*_*s*_*/*2 bases upstream and *W*_*s*_*/*2 bases downstream of the cytosine) chosen uniformly at random among all possible cytosines that belonged to that particular context (either CG, CHH or CHG), in which *n/*2 were methylated (i.e., have a methylation level of at least 0.5) and *n/*2 were unmethylated (i.e., have a methylation level below 0.5). The reason to balance methylated/unmethylated examples is explained in Section 2. In some cases, *n* was limited by the number of available methylated cytosine genome-wide (e.g., CHH in Arabidopsis, see Supplemental Table 4). Even in those cases, however, we kept the training set balanced in terms of methylated/unmethylated cytosine. For the combined context, the training set consisted of an equal number of examples for CG, CHG and CHH. The reason to balance the contexts is explained in Section 2. Since the DNA sequence were one-hot encoded, the training set was composed of *n* binary matrices of size *W*_*s*_ × 4. Several choices of the window size *W*_*s*_ and the training set size were tested, as explained in Section 2.3.

For classifiers that rely on genomic annotations (in addition to the primary DNA sequence) the one-hot encoded *W*_*s*_ × 4 input was augmented with a few bit-vectors representing the annotations. We used two bit-vectors for each functional element, one for each strand. The binary values of these bit-vectors indicated the annotation status of each nucleotide in the window. If a nucleotide was contained in a particular functional element (e.g., coding sequence), the corresponding value in the strand-specific bit-vector was one (and zero otherwise). Supplementary Table 5 shows the functional elements used for each species.

For classifiers that rely on the methylation level of neighboring cytosines, the context-specific training set was composed by *n* vectors of length *W*_*p*_, where the first *W*_*p*_*/*2 components of the vector are methylation levels (in the range [0, 1]) of the cytosines upstream and the second *W*_*p*_*/*2 components of the vector are methylation levels (in the range [0, 1]) of cytosines downstream of a cytosine chosen uniformly at random among all possible cytosines that belong to that particular context (either CG, CHH or CHG). For the combined context, the training set was composed by an equal number of examples from CG, CHG and CHH. Again, we made sure that the training set was balanced: *n/*2 samples had a center cytosine which was methylated, and *n/*2 samples had an unmethylated center cytosine. Please note that while the center cytosine is context-specific, the vector contained methylation levels for cytosines in any context, as long they had sufficient read coverage (i.e., more than ten reads).

In all experiments eighty percent of the data was used for training, ten percent was used for validation, and ten percent was used for testing. Validation and test data sets had the same characteristics of the training set, but we made sure no DNA sequence in the training set appeared in the test set.

### Classifiers

We first studied the prediction accuracy of a simple Random Forest classifier (RF) because RF has been used in the literature for this problem (see, e.g., [37, 38]). The Random Forest classifier was implemented using Python Scikit-learn (version 0.24.2) and trained with fifty estimators and unlimited trees depth.

The most effective ML methods in the literature to predict cytosine methylation are, however, based on deep learning (see, e.g., [32, 33, 34, 35, 36]). To carry out an extensive set of prediction experiments, we created a simple deep learning architecture based on CNNs. The model was kept simple (i.e., just a few hidden layers) to avoid over-fitting. For the same reason we purposely did not optimize the hyper-parameters of AMPS. Doing so would allow the network to achieve a higher accuracy on specific data sets, but possibly hamper its generalization abilities.

We called our architecture for the prediction of cytosine methylation <u>A</u>nnotation-based <u>M</u>ethylation <u>P</u>rediction from <u>S</u>equence (AMPS). As explained in the previous section, the input to AMPS is a matrix of the size of *W*_*s*_×(4+*a*) where *a* is the number of bit-vectors representing the annotations (*a* = 0 when AMPS only uses the sequence, i.e., no annotations). The input was first processed by a 1D convolutional layer with kernel size of (4 + *a*). This convolution layer had 16 channels followed by a ReLU function. The next layer was a fully connected layer with 128 nodes using a ReLU activation function. To avoid over fitting, a dropout ratio of 0.5 was used for the fully connected layer. The last layer was a single node using a sigmoid activation function. A stochastic gradient descent optimizer was used, and the loss function was binary cross-entropy.

The input to the network for the prediction of cytosine methylation from neighboring cytosines was a vector of methylation levels in the range [0, 1] of length *W*_*p*_. The network was composed of four fully connected layers with 20, 16, 8, and one node, respectively. The hidden layers used ReLU as their activation function. A dropout ratio of 0.5 was used to prevent over fitting. A stochastic gradient descent was used for optimization, and binary cross-entropy was used for the loss function.

## 4 Discussion and Conclusions

In this study, we investigated the problem of predicting cytosine methylation in plants from either the DNA sequence or the neighboring cytosines. To the best of our knowledge, this is the first time that independent predictions for different contexts and plant species have been carried out and compared. We can summarize our findings in three major categories.

Our first finding is that the cytosine methylation prediction from the sequence is more accurate when a context-specific species-specific classifier is used. Combining the contexts during training, which is what most paper in the literature have done so far (although some only focus on CG), degrades the performance of the classifier. Our study suggests that context-specific species-specific predictive models are necessary for obtaining the best overall predictive performance for cytosine methylation from the primary sequence in plants, and possibly elsewhere.

The second finding is that the predictive accuracy of cytosine methylation from the methylation levels of neighboring cytosines is higher than the predictive accuracy obtained from the sequence only (with or without annotation). This is consistent with results reported in the literature for other organisms (mostly vertebrates). However, to the best of our knowledge, no study has compared predictions from neighboring cytosine across multiple organisms or across contexts. In fact, cross-species and cross-context prediction appears sufficiently accurate, which opens the possibility of methylation imputation across species. Interestingly, while the easiest context to predict from the sequence is CHG, the easiest context to predict from neighboring methylation levels is CG.

The final, and perhaps most intriguing, finding of our study is that using genome annotation data dramatically improves the predictive accuracy of cytosine methylation from the sequence. Our experimental results show that our AMPS classifier with annotation has a much higher accuracy than AMPS without annotations. In addition, AMPS with annotations performs better than CpGenie on most of the context/organism pairs. Since CpGenie has a more complex/optimized network than AMPS, one can speculate that extending the CpGenie architecture by taking advantage of genome annotations could produce a classifier that is better than CpGenie or AMPS on all context and species.

A potential direction of improvement in this line of research is to provide additional functional annotations in input to the classifier. For instance, it is known that transponsable elements are silenced in plants via DNA methylation (see, e.g., [44]), so it is possible that repeat annotations would further improve AMPS prediction accuracy from the sequence.

## Acknowledgements

The authors want to thank Felix Krueger (the author of Bismark) for helpful discussions on DNA methylation analysis pipeline.

## Abbreviations

AMPS: Annotation-based Methylation Prediction from Sequence (classifier)
RF: random forest (classifier)
CNN: convolutional neural network
WGBS: Whole genome bisulfite sequencing
BS-Seq: Sodium bisulfite sequencing
SRA: Sequence Read Archive

## Availability of data and materials

Whole Genome Bisulfite Sequencing data supporting the conclusions of this article are available in the NCBI SRA repository with accession numbers SRR3171614, SRR618545, SRR618546, SRR618547, SRR503393 and SRR5430777 for Arabidopsis, rice, cucumber and tomato, respectively. Cowpea BS-Seq data were deposited in the European Nucleotide Archive with accession number PRJEB52355. Code and scripts are available in the public Github repository https://github.com/ucrbioinfo/AMPS

## Authors’ contributions

SS and SL conceived the project and designed the experiments. SL supervised the project. SS developed AMPS, and carried out all the experiments. DF and MO conceived the cowpea trials to target differential epigenetic modifications and carried out the BS-Seq experiments. SS and SL wrote the manuscript. DF and MO suggested edits and changes to the manuscript.

## References

[1] Schübeler, D.: Function and information content of DNA methylation. Nature 517(7534), 321–326 (2015)

[2] Yang, X., Han, H., De Carvalho, D.D., Lay, F.D., Jones, P.A., Liang, G.: Gene body methylation can alter gene expression and is a therapeutic target in cancer. Cancer cell 26(4), 577–590 (2014)

[3] Seymour, D.K., Gaut, B.S.: Phylogenetic shifts in gene body methylation correlate with gene expression and reflect trait conservation. Molecular Biology and Evolution 37(1), 31–43 (2020)

[4] Vinson, C., Chatterjee, R.: CG methylation. Epigenomics 4(6), 655–663 (2012)

[5] Jeziorska, D.M., Murray, R.J., De Gobbi, M., Gaentzsch, R., Garrick, D., Ayyub, H., Chen, T., Li, E., Telenius, J., Lynch, M., et al.: DNA methylation of intragenic CpG islands depends on their transcriptional activity during differentiation and disease. Proceedings of the National Academy of Sciences 114(36), 7526–7535 (2017)

[6] Straussman, R., Nejman, D., Roberts, D., Steinfeld, I., Blum, B., Benvenisty, N., Simon, I., Yakhini, Z., Cedar, H.: Developmental programming of CpG island methylation profiles in the human genome. Nature structural & molecular biology 16(5), 564–571 (2009)

[7] Moore, L.D., Le, T., Fan, G.: DNA methylation and its basic function. Neuropsychopharmacology 38(1), 23–38 (2013)

[8] Aceituno, F.F., Moseyko, N., Rhee, S.Y., Gutiérrez, R.A.: The rules of gene expression in plants: organ identity and gene body methylation are key factors for regulation of gene expression in Arabidopsis thaliana. BMC genomics 9(1), 1–14 (2008)

[9] Doi, A., Park, I.-H., Wen, B., Murakami, P., Aryee, M.J., Irizarry, R., Herb, B., Ladd-Acosta, C., Rho, J., Loewer, S., et al.: Differential methylation of tissue-and cancer-specific CpG island shores distinguishes human induced pluripotent stem cells, embryonic stem cells and fibroblasts. Nature genetics 41(12), 1350–1353 (2009)

[10] Das, P.M., Singal, R.: DNA methylation and cancer. Journal of clinical oncology 22(22), 4632–4642 (2004)

[11] Mill, J., Tang, T., Kaminsky, Z., Khare, T., Yazdanpanah, S., Bouchard, L., Jia, P., Assadzadeh, A., Flanagan, J., Schumacher, A., et al.: Epigenomic profiling reveals DNA-methylation changes associated with major psychosis. The American Journal of Human Genetics 82(3), 696–711 (2008)

[12] Apazoglou, K., Adouan, W., Aubry, J.-M., Dayer, A., Aybek, S.: Increased methylation of the oxytocin receptor gene in motor functional neurological disorder: a preliminary study. Journal of Neurology, Neurosurgery & Psychiatry 89(5), 552–554 (2018)

[13] Zhang, X., Yazaki, J., Sundaresan, A., Cokus, S., Chan, S.W.-L., Chen, H., Henderson, I.R., Shinn, P., Pellegrini, M., Jacobsen, S.E., et al.: Genome-wide high-resolution mapping and functional analysis of DNA methylation in Arabidopsis. Cell 126(6), 1189–1201 (2006)

[14] Lister, R., O’Malley, R.C., Tonti-Filippini, J., Gregory, B.D., Berry, C.C., Millar, A.H., Ecker, J.R.: Highly integrated single-base resolution maps of the epigenome in Arabidopsis. Cell 133(3), 523–536 (2008)

[15] Harris, K.D., Zemach, A.: Contiguous and stochastic CHH methylation patterns of plant DRM2 and CMT2 revealed by single-read methylome analysis. Genome biology 21(1), 1–19 (2020)

[16] Kenchanmane Raju, S.K., Ritter, E.J., Niederhuth, C.E.: Establishment, maintenance, and biological roles of non-CG methylation in plants. Essays in biochemistry 63(6), 743–755 (2019)

[17] To, T.K., Yamasaki, C., Oda, S., Tominaga, S., Kobayashi, A., Tarutani, Y., Kakutani, T.: Local and global crosstalk among heterochromatin marks drives DNA methylome patterning in Arabidopsis. Nature Communications 13(1), 1–10 (2022)

[18] de Mendoza, A., Poppe, D., Buckberry, S., Pflueger, J., Albertin, C.B., Daish, T., Bertrand, S., de la Calle-Mustienes, E., Gómez-Skarmeta, J.L., Nery, J.R., et al.: The emergence of the brain non-CpG methylation system in vertebrates. Nature ecology & evolution 5(3), 369–378 (2021)

[19] Kozlenkov, A., Li, J., Apontes, P., Hurd, Y.L., Byne, W.M., Koonin, E.V., Wegner, M., Mukamel, E.A., Dracheva, S.: A unique role for DNA (hydroxy) methylation in epigenetic regulation of human inhibitory neurons. Science advances 4(9), 6190 (2018)

[20] He, Y., Ecker, J.R.: Non-CG methylation in the human genome. Annual review of genomics and human genetics 16, 55–77 (2015)

[21] Cui, D., Xu, X.: DNA methyltransferases, DNA methylation, and age-associated cognitive function. International journal of molecular sciences 19(5), 1315 (2018)

[22] Perzel Mandell, K.A., Eagles, N.J., Wilton, R., Price, A.J., Semick, S.A., Collado-Torres, L., Ulrich, W.S., Tao, R., Han, S., Szalay, A.S., et al.: Genome-wide sequencing-based identification of methylation quantitative trait loci and their role in schizophrenia risk. Nature communications 12(1), 1–12 (2021)

[23] Tan, F., Zhou, C., Zhou, Q., Zhou, S., Yang, W., Zhao, Y., Li, G., Zhou, D.-X.: Analysis of chromatin regulators reveals specific features of rice DNA methylation pathways. Plant physiology 171(3), 2041–2054 (2016)

[24] Bewick, A.J., Schmitz, R.J.: Gene body DNA methylation in plants. Current opinion in plant biology 36, 103–110 (2017)

[25] Wang, H., Beyene, G., Zhai, J., Feng, S., Fahlgren, N., Taylor, N.J., Bart, R., Carrington, J.C., Jacobsen, S.E., Ausin, I.: CG gene body DNA methylation changes and evolution of duplicated genes in cassava. Proceedings of the National Academy of Sciences 112(44), 13729–13734 (2015). doi:10.1073/pnas.1519067112. https://www.pnas.org/doi/pdf/10.1073/pnas.1519067112

[26] Bewick, A.J., Ji, L., Niederhuth, C.E., Willing, E.-M., Hofmeister, B.T., Shi, X., Wang, L., Lu, Z., Rohr, N.A., Hartwig, B., Kiefer, C., Deal, R.B., Schmutz, J., Grimwood, J., Stroud, H., Jacobsen, S.E., Schneeberger, K., Zhang, X., Schmitz, R.J.: On the origin and evolutionary consequences of gene body DNA methylation. Proceedings of the National Academy of Sciences 113(32), 9111–9116 (2016). doi:10.1073/pnas.1604666113. https://www.pnas.org/doi/pdf/10.1073/pnas.1604666113

[27] Niederhuth, C.E., Bewick, A.J., Ji, L., Alabady, M.S., Do Kim, K., Li, Q., Rohr, N.A., Rambani, A., Burke, J.M., Udall, J.A., et al.: Widespread natural variation of DNA methylation within angiosperms. Genome biology 17(1), 1–19 (2016)

[28] Jain, M., Olsen, H.E., Paten, B., Akeson, M.: The Oxford Nanopore MinION: delivery of nanopore sequencing to the genomics community. Genome biology 17(1), 1–11 (2016)

[29] Krueger, F., Andrews, S.R.: Bismark: a flexible aligner and methylation caller for bisulfite-seq applications. bioinformatics 27(11), 1571–1572 (2011)

[30] Chen, P.-Y., Cokus, S.J., Pellegrini, M.: BS Seeker: precise mapping for bisulfite sequencing. BMC bioinformatics 11(1), 1–6 (2010)

[31] Harris, E.Y., Ounit, R., Lonardi, S.: BRAT-nova: fast and accurate mapping of bisulfite-treated reads. Bioinformatics 32(17), 2696–2698 (2016)

[32] Angermueller, C., Lee, H.J., Reik, W., Stegle, O.: DeepCpG: accurate prediction of single-cell DNA methylation states using deep learning. Genome biology 18(1), 1–13 (2017)

[33] Li, R.A., Liu, Z.: A hybrid deep neural network for robust single-cell genome-wide DNA methylation detection. In: Proceedings of the 12th ACM Conference on Bioinformatics, Computational Biology, and Health Informatics, pp. 1–6 (2021)

[34] Tian, Q., Zou, J., Tang, J., Fang, Y., Yu, Z., Fan, S.: MRCNN: a deep learning model for regression of genome-wide DNA methylation. BMC genomics 20(2), 1–10 (2019)

[35] Zeng, H., Gifford, D.K.: Predicting the impact of non-coding variants on DNA methylation. Nucleic acids research 45(11), 99–99 (2017)

[36] De Waele, G., Clauwaert, J., Menschaert, G., Waegeman, W.: CpG transformer for imputation of single-cell methylomes. Bioinformatics 38(3), 597–603 (2022)

[37] Zhang, W., Spector, T.D., Deloukas, P., Bell, J.T., Engelhardt, B.E.: Predicting genome-wide DNA methylation using methylation marks, genomic position, and DNA regulatory elements. Genome biology 16(1), 1–20 (2015)

[38] Zheng, Y., Joyce, B.T., Liu, L., Zhang, Z., Kibbe, W.A., Zhang, W., Hou, L.: Prediction of genome-wide DNA methylation in repetitive elements. Nucleic acids research 45(15), 8697–8711 (2017)

[39] Du, J., Johnson, L.M., Jacobsen, S.E., Patel, D.J.: DNA methylation pathways and their crosstalk with histone methylation. Nature Reviews Molecular Cell Biology 16(9), 519–532 (2015). doi:10.1038/nrm4043

[40] Omirou, M., Ioannides, I.M., Fasoula, D.A.: Optimizing resource allocation in a cowpea (Vigna unguiculata L. Walp.) landrace through whole-plant field phenotyping and non-stop selection to sustain increased genetic gain across a decade. Frontiers in Plant Science 10 (2019). doi:10.3389/fpls.2019.00949

[41] Lonardi, S., Muñoz-Amatriaín, M., Liang, Q., Shu, S., Wanamaker, S.I., Lo, S., Tanskanen, J., Schulman, A.H., Zhu, T., Luo, M.-C., Alhakami, H., Ounit, R., Hasan, A.M., Verdier, J., Roberts, P.A., Santos, J.R.P., Ndeve, A., Doležel, J., Vrána, J., Hokin, S.A., Farmer, A.D., Cannon, S.B., Close, T.J.: The genome of cowpea (Vigna unguiculata [L.] Walp.). The Plant Journal 98(5), 767–782 (2019). doi:10.1111/tpj.14349. https://onlinelibrary.wiley.com/doi/pdf/10.1111/tpj.14349

[42] Liu, Y.-Y., Yang, K.-Z., Wei, X.-X., Wang, X.-Q.: Revisiting the phosphatidylethanolamine-binding protein (pebp) gene family reveals cryptic flowering locus t gene homologs in gymnosperms and sheds new light on functional evolution. New Phytologist 212(3), 730–744 (2016). doi:10.1111/nph.14066. https://nph.onlinelibrary.wiley.com/doi/pdf/10.1111/nph.14066

[43] Bolger, A.M., Lohse, M., Usadel, B.: Trimmomatic: a flexible trimmer for Illumina sequence data. Bioinformatics 30(15), 2114–2120 (2014)

[44] Sigman, M.J., Slotkin, R.K.: The first rule of plant transposable element silencing: Location, location, location. The Plant cell 28(2), 304–313 (2016). doi:10.1105/tpc.15.00869

