## Supplemental material for "On the Prediction of non-CG DNA Methylation"

### On the Prediction of non-CG DNA Methylation
Supplementary material

Saleh Sereshki, Michalis Omirou, Dionysia Fasoula and Stefano Lonardi

#### Supplementary Tables

| species | source | assembly version | size (Mb) |
|---|---|---|---|
| Arabidopsis | The Arabidopsis Information Resource | TAIR10.1 | 119.6 |
| cowpea | JGI Phytozome | IT97K-499-35 v.1 | 519.4 |
| rice | International Rice Genome Sequencing Project | IRGSP-1.0 | 373.2 |
| cucumber | The Cucumber Genome Initiative | GCF_000004075.3_9930_V3 | 226.6 |
| tomato | Solanaceae Genomics Project | GCF_000188115.4_SL3.0 | 828.3 |

Supplementary Table 1: Source, version and genome size of the genome assemblies used in this study

| species | SRA source | number of reads | % of reads mapped | read length (after trimming) | read type |
|---|---|---|---|---|---|
| Arabidopsis | SRR3171614 | 209,561,030 | 54.40% | 50 | single end |
| cowpea | PRJEB52355 | 141,137,614 | 39.30% | 150 | paired end |
| rice | SRR618545-7 | 622,913,368 | 47.30% | 50 (37) | single end |
| cucumber | SRR5430777 | 59,999,999 | 50.10% | 90 | paired end |
| tomato | SRR503393 | 84,127,751 | 77.70% | 101 (91) | paired end |

Supplementary Table 2: Summary of the BS-Seq data sets used in this study: SRA/EMBL source, number of reads, read length (before and after trimming), read type and % of reads uniquely mapped

| species | genome size (Mb) | average read coverage | average cytosine coverage |
|---|---|---|---|
| Arabidopsis | 119.6 | 48x | 21x |
| cowpea | 519.4 | 32x | 12x |
| rice | 373.2 | 29x | 12x |
| cucumber | 226.6 | 24x | 9x |
| tomato | 828.3 | 16x | 5x |

Supplementary Table 3: Genome sizes, average genome coverage from Bismark mapped reads, average cytosine coverage from Bismark mapped reads

| | | methylated cytosine (%) | | | |
|---|---|---|---|---|---|
| species | Cs with sufficient coverage (%) | CG | CHG | CHH | ALL |
| Arabidopsis | 62.29% | 27.54% | 7.72% | 0.68% | 6.00% |
| cowpea | 44.97% | 60.40% | 47.54% | 4.41% | 16.30% |
| rice | 37.87% | 54.15% | 26.87% | 2.66% | 18.21% |
| cucumber | 22.63% | 56.58% | 24.93% | 5.22% | 15.05% |
| tomato | 9.13% | 89.56% | 62.88% | 2.73% | 19.01% |

Supplementary Table 4: Summary of cytosine methylation statistics for the species in this study; the second column shows the percentage of cytosines that have a coverage of more than ten reads after mapping BS-Seq reads with Bismark; the rest of the columns show the percentage of methylated cytosines in each individual context (only for the cytosines that had sufficient coverage)

| species | number of annotated elements |
|---|---|
| Arabidopsis | genes (38,310), exons (324,656), CDS (286,173) |
| cowpea | genes (31,947), CDS (325,882) |
| rice | mRNA (44,784), exons (198,572) |
| cucumber | gene (23,636), exons (254,378), CDS (201,484) |
| tomato | genes (30,017), exons (310,488), CDS (235,394) |

Supplementary Table 5: Number of available annotations in each species

| | methylated | | | | unmethylated | | | |
|---|---|---|---|---|---|---|---|---|
| species | CG (%) | CHG (%) | CHH (%) | ALL (%) | CG (%) | CHG (%) | CHH (%) | ALL (%) |
| Arabidopsis | 72.61 | 20.57 | 6.82 | 100 | 11.85 | 16.26 | 71.88 | 100 |
| cowpea | 40.79 | 40.23 | 18.97 | 100 | 5.23 | 8.76 | 86.01 | 100 |
| rice | 63.91 | 27.95 | 8.13 | 100 | 11.88 | 17.05 | 71.07 | 100 |
| cucumber | 52.59 | 24.28 | 23.13 | 100 | 6.93 | 12.82 | 80.25 | 100 |
| tomato | 51.15 | 39.11 | 9.74 | 100 | 1.40 | 5.36 | 93.25 | 100 |

Supplementary Table 6: Distribution of methylated and unmethylated cytosines for each individual context

### Supplementary Figures

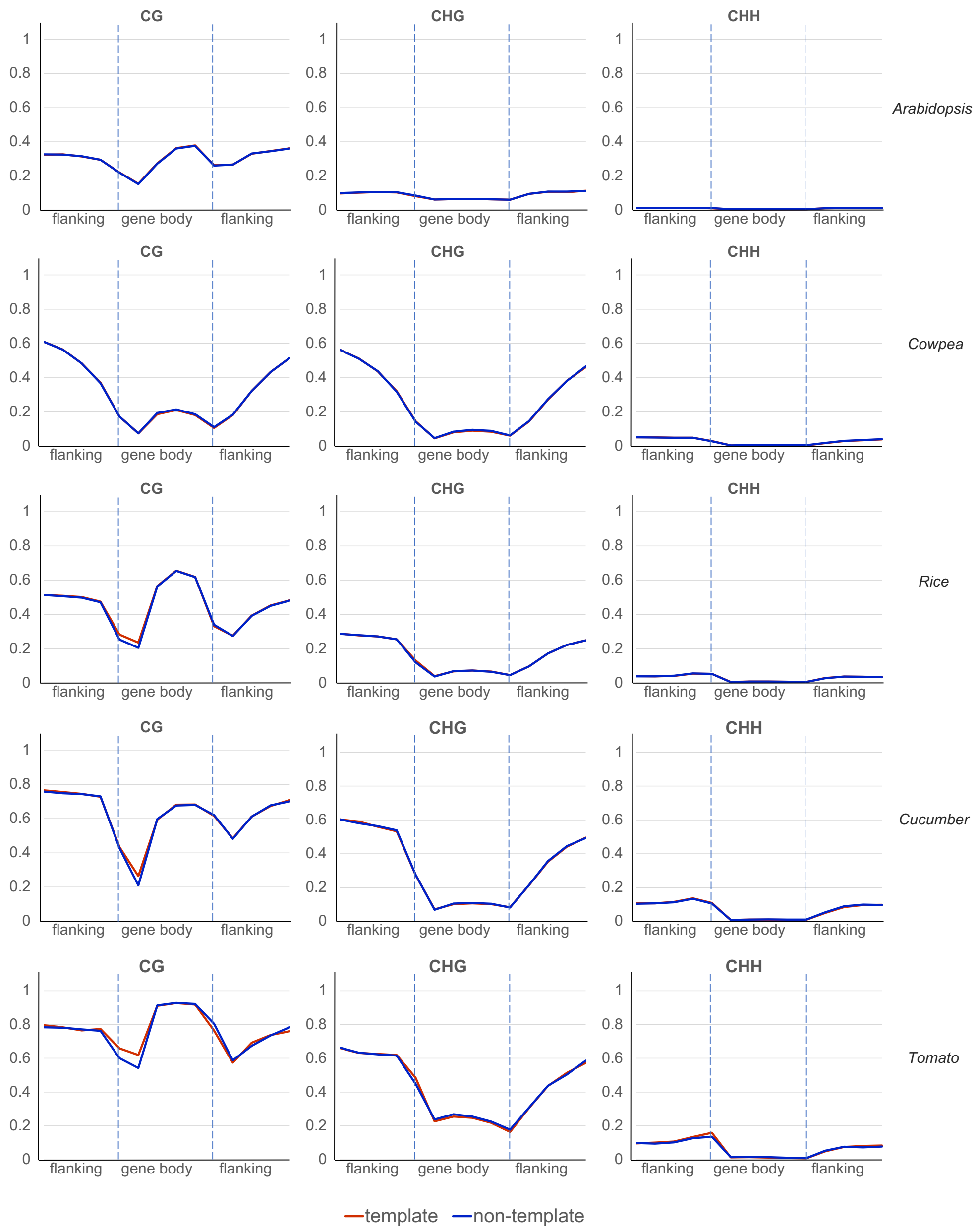


Supplementary Figure 1: Context-specific, species-specific gene body methylation levels in gene bodies and 2000 bp flanking regions (upstream and downstream) for template and non-template strands, when averaged over all genes

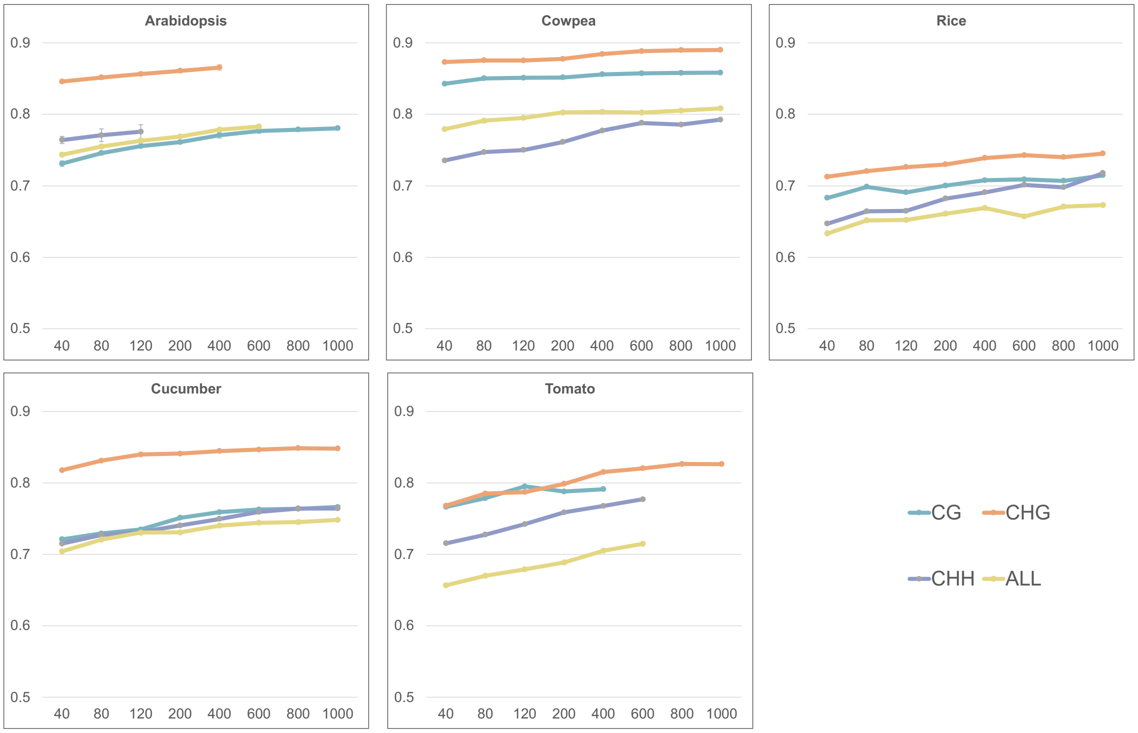


Supplementary Figure 2: Prediction accuracy as a function of the training set size for AMPS with annotations (the x-axis represents thousands of samples); on the Arabidopsis plot, the error bars indicate the standard deviation of the accuracy over six independent experiments.

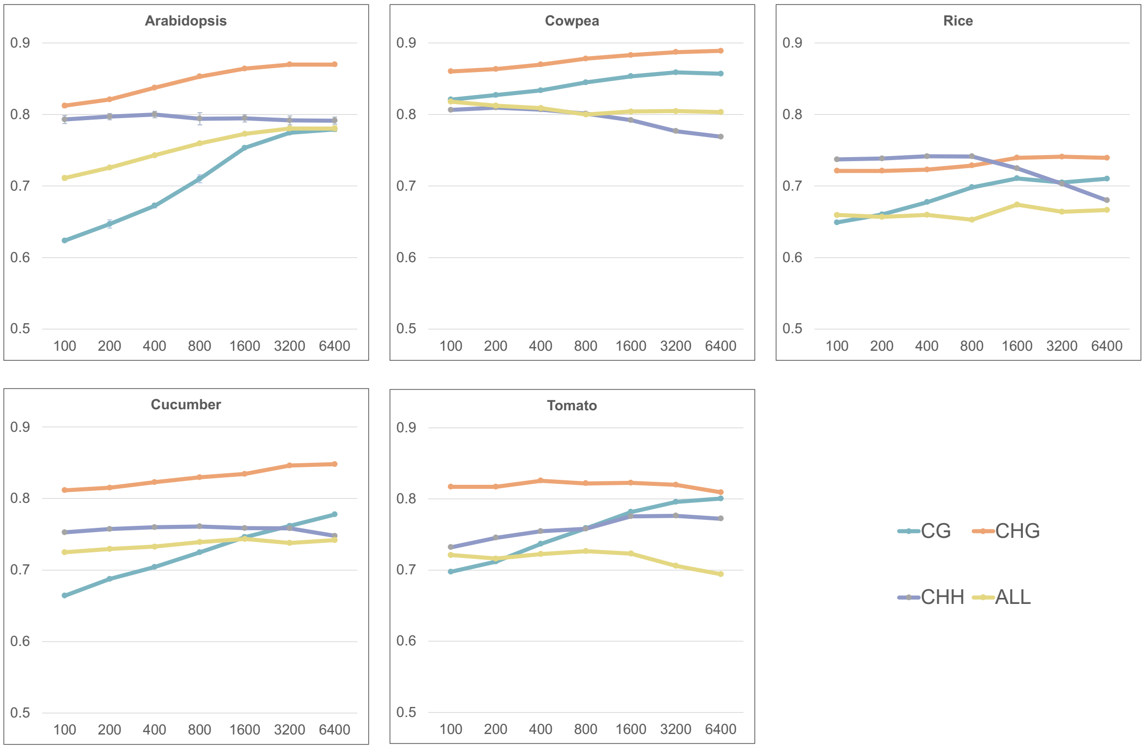


Supplementary Figure 3: Prediction accuracy as a function of the window size for AMPS with annotations (the x-axis represents the window size in base pairs); on the Arabidopsis plot, the error bars indicate the standard deviation of the accuracy over six independent experiments.

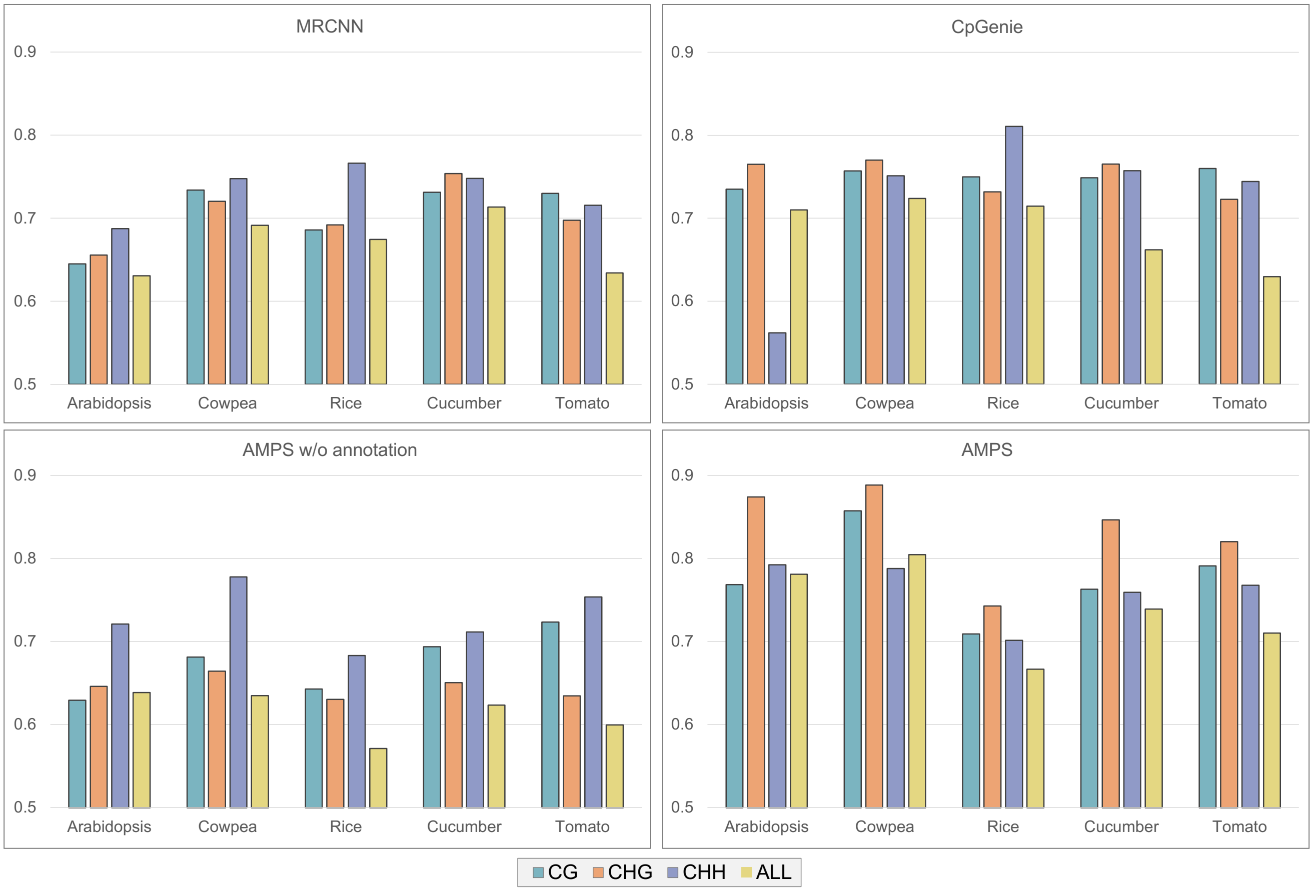


Supplementary Figure 4: Context-specific species-specific cross-validation accuracy for methylation prediction from DNA sequence for MRCNN, CPGenie, AMPS without annotations and AMPS with annotations

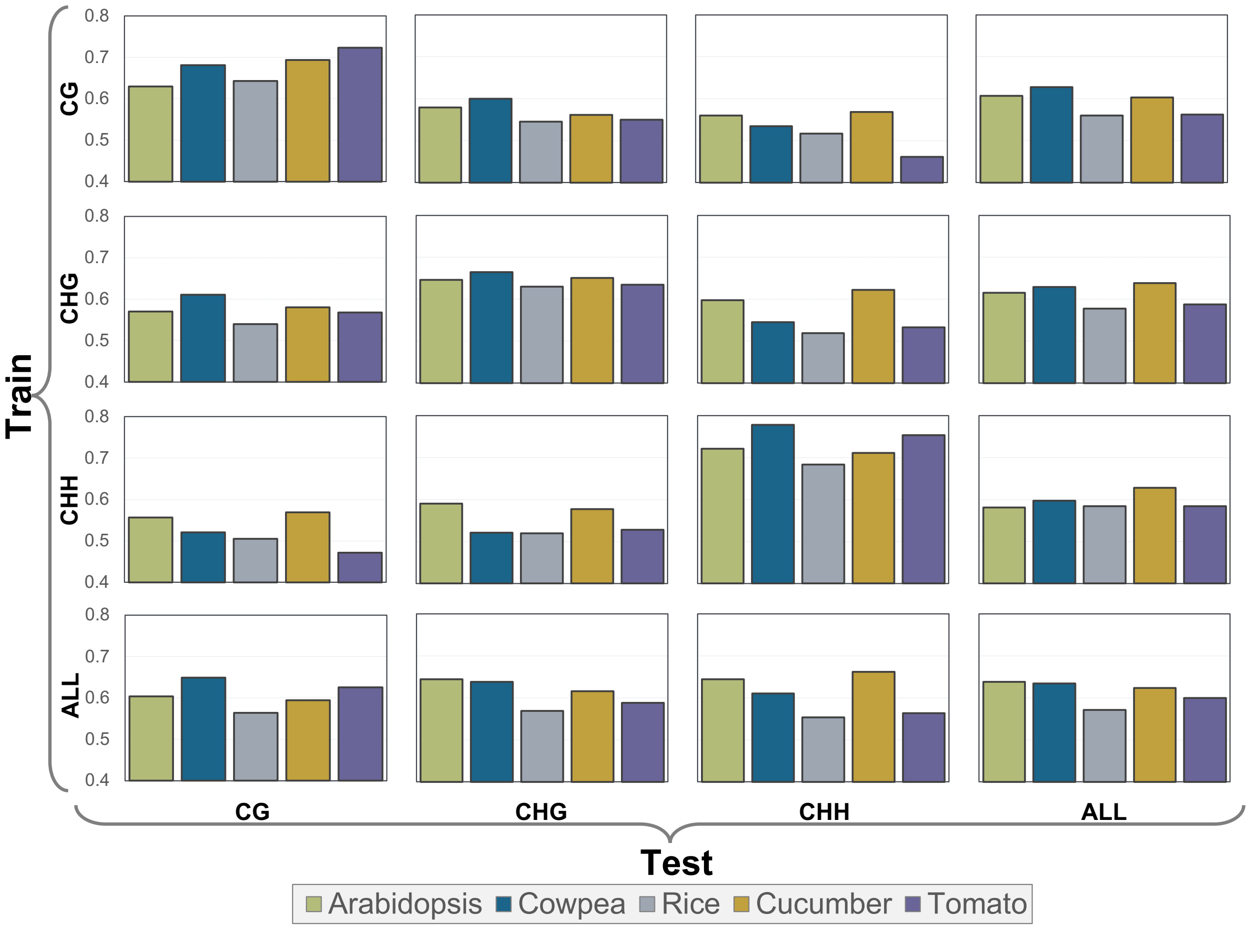


Supplementary Figure 5: Cross-context methylation prediction for AMPS without annotation

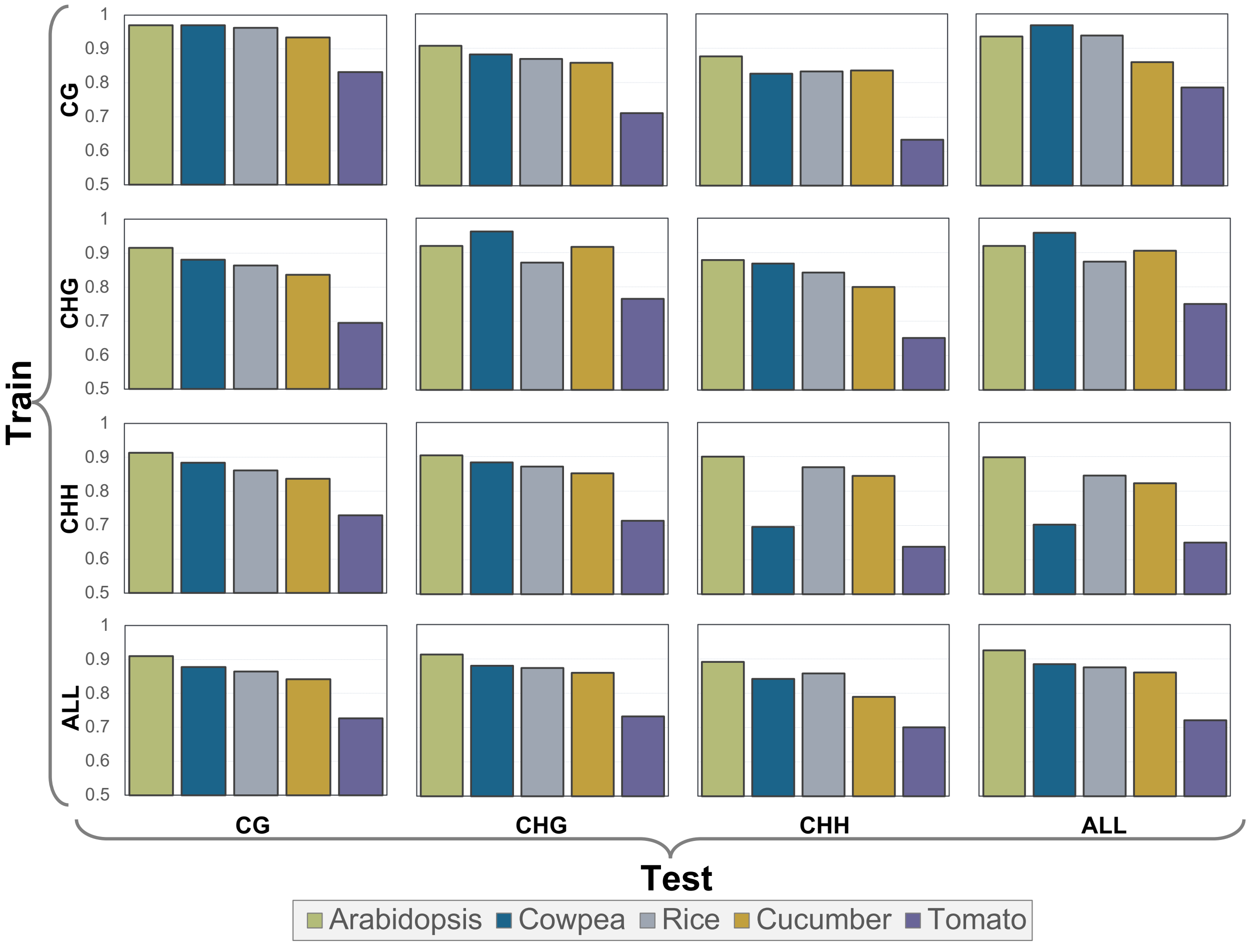


Supplementary Figure 6: Cross-context methylation prediction from neighboring cytosines

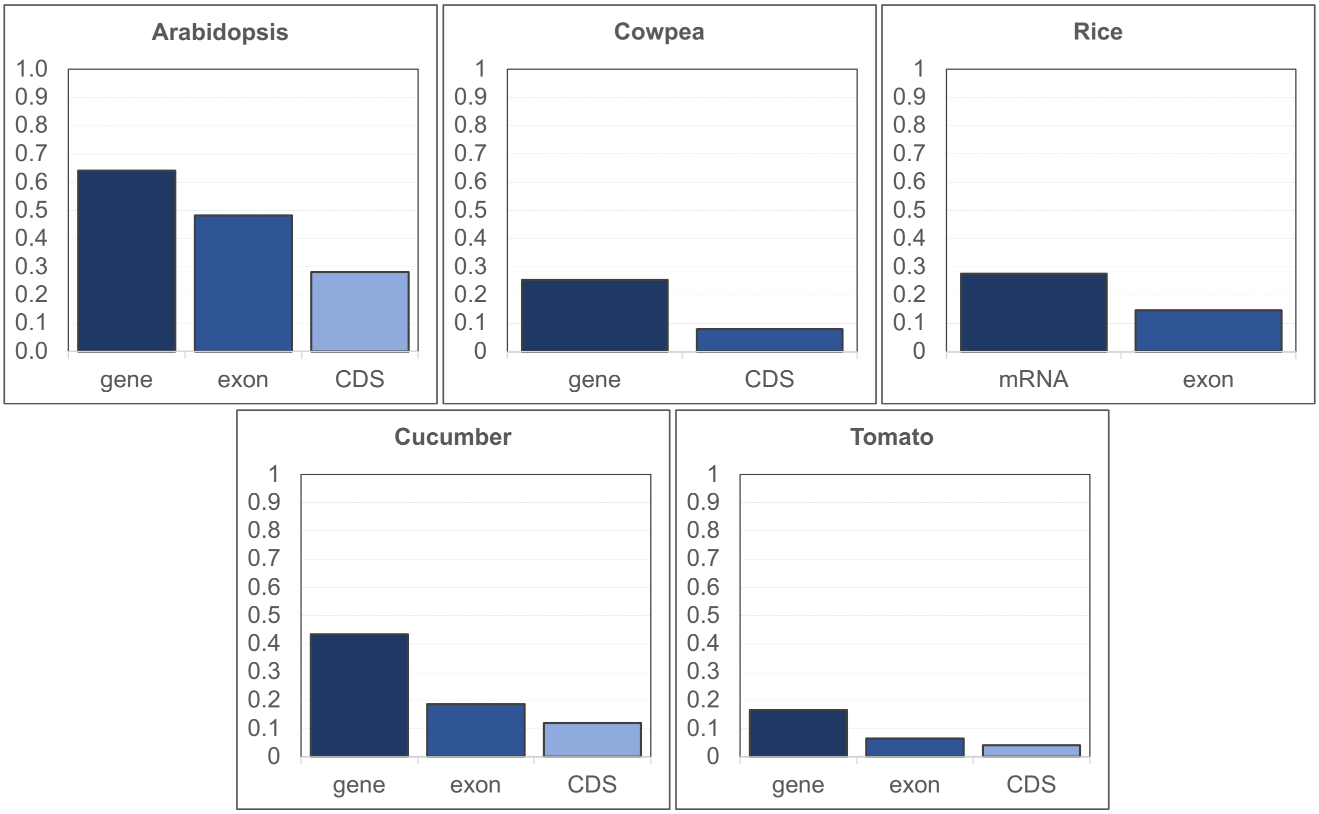


Supplementary Figure 7: Fraction of each genome covered by a functional annotation (see Supplemental Table 5 for the list)

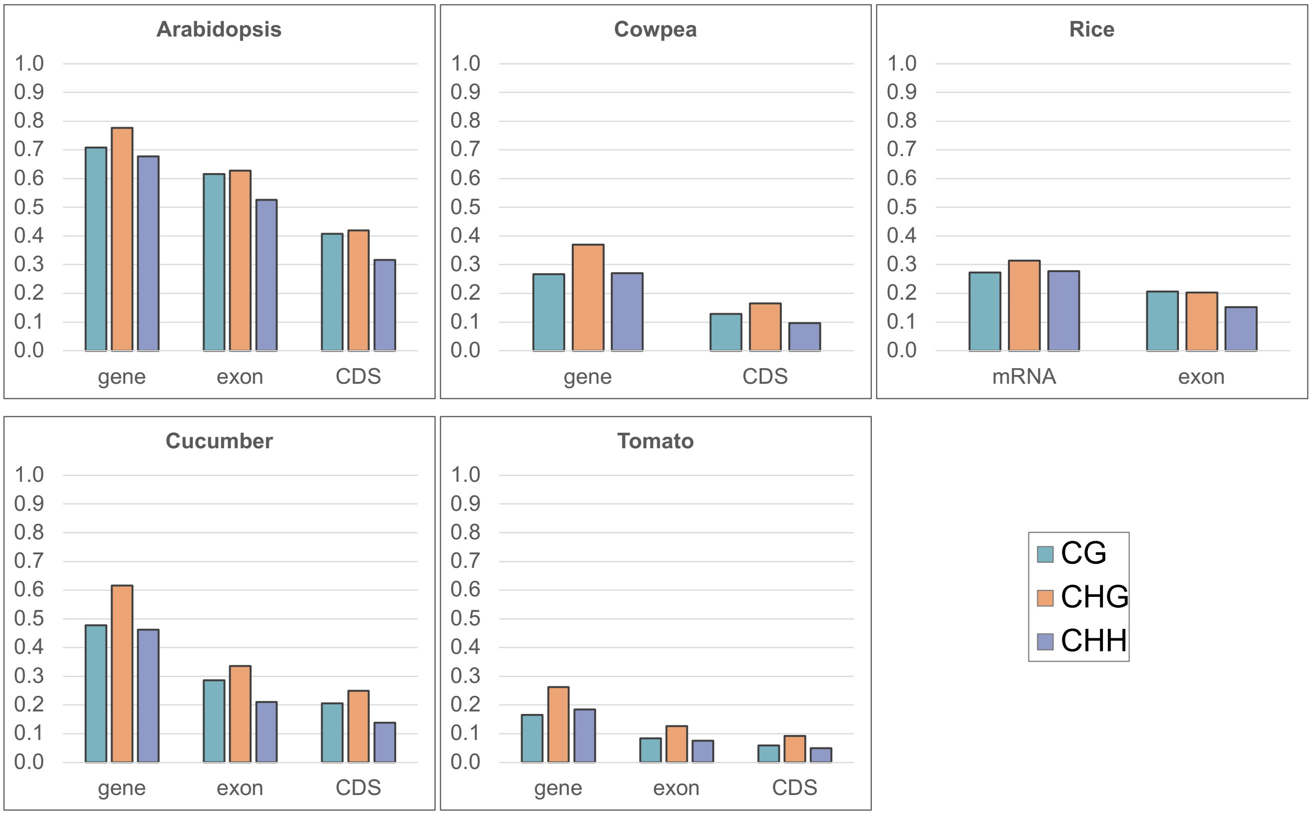


Supplementary Figure 8: Context-specific species-specific fraction of all cytosines covered by a functional annotation (see Supplemental Table 5 for the list)

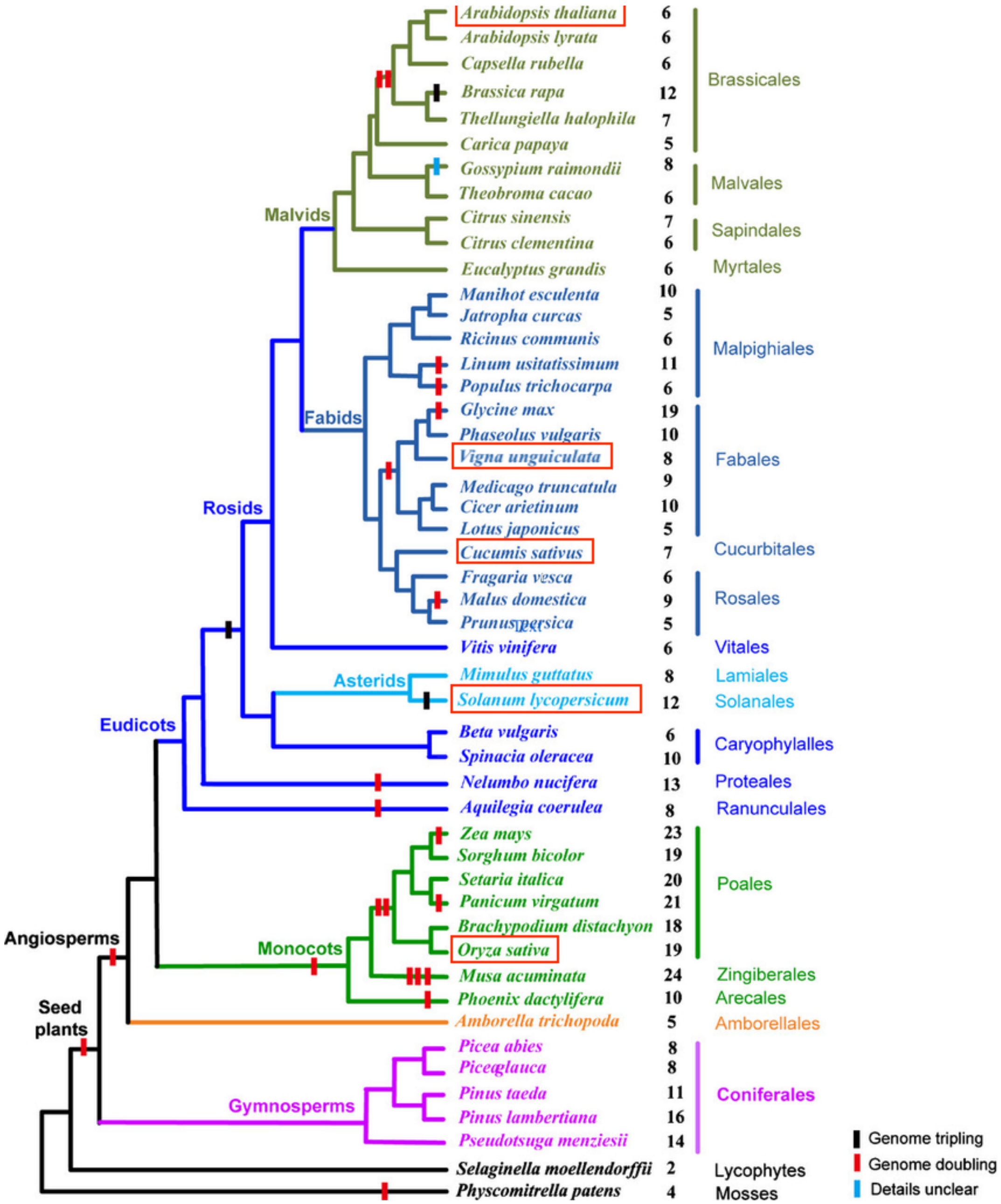


Supplementary Figure 9: A phylogenetic tree of land plants, decorated with whole-genome duplication events (adapted from doi:10.1111/nph.14066); the five species included in this study are hightlighted in red
